# Live-cell co-translational folding tracking reveals bidirectional coupling between translation and folding

**DOI:** 10.64898/2026.07.12.738044

**Authors:** Rhiannon M. Sears, Luis U. Aguilera, Tristan M. Bunting, Ning Zhao

**Author notes:** These authors contributed equally to this work (Rhiannon M. Sears, Luis U. Aguilera).

## Abstract

Translation elongation and protein folding have long been proposed to coordinate during co-translational folding, yet the lack of technologies capable of simultaneously tracking both processes in live cells has hindered mechanistic understanding of this relationship. Here, we developed co-translational folding tracking (coTFT), a live-cell imaging platform that directly and simultaneously tracks translation and folding from individual mRNAs. Using reporters with distinct folding kinetics, we found that differences in folding kinetics were accompanied by corresponding changes in translation elongation rates. Conversely, altering translation elongation markedly affected protein folding outcomes. Combining coTFT with mathematical modeling enabled estimation of reporter folding times on translating ribosomes in live cells, confirming their distinct folding kinetics. Together, our results reveal that translation elongation and folding are bidirectionally coupled during co-translational folding.

---

Protein folding can begin while nascent chains are still being synthesized by ribosomes, a process known as co-translational folding (coF) (*1–5*). By coupling protein folding to the vectorial emergence of nascent chains from the ribosome, coF is thought to enhance protein folding efficiency (*5–7*). Translation elongation has long been proposed to regulate coF by controlling the timing of nascent chain emergence (*8–12*). Conversely, folding of nascent chains can generate mechanical pulling forces that feed back on the ribosome and potentially influence translation elongation (*13*). Despite decades of study, coF has been investigated primarily using in vitro systems (*12*, *13*) or ensemble measurements (*14*, *15*). Moreover, most existing assays infer folding from the function of mature proteins (*16*), providing indirect, endpoint readouts that cannot capture folding as it occurs on translating ribosomes. As a result, whether and how translation elongation and protein folding are coordinated during coF in live cells remains unresolved, limiting mechanistic insight into coF regulation.

To address this question, we developed co-translational folding tracking (coTFT), a live-cell imaging platform that enables direct and simultaneous tracking of translation and folding from individual mRNAs in their native intracellular environment. Applying coTFT to reporters with distinct folding kinetics reveals a novel regulatory principle in which translation elongation and protein folding are bidirectionally coupled to coordinate co-translational protein biogenesis in live cells.

## Tracking single-mRNA co-translational folding in live cells

In this study, we developed a coTFT technology that enables direct visualization of protein folding on translating ribosomes from individual mRNAs in live cells. As illustrated in **Fig. 1A**, the coTFT reporter consists of an N-terminal spaghetti monster HA (smHA) tag containing 10 HA epitopes embedded within an mRuby scaffold (*17*), followed by tandem repeats of dark GFP (dGFP) (*18*). Upon translation initiation, a co-expressed anti-HA intrabody (ib) (*19*) fused to HaloTag-JF646 (anti-HA-ib-Halo) binds the smHA tag and labels nascent chains (**Fig. 1A**, magenta). As dGFP folds, a co-expressed conformation-sensitive (CS) anti-GFP intrabody fused to tandem mStayGold (mSG, anti-GFP-ib-mSG) selectively binds folded dGFP, marking nascent chains undergoing folding (**Fig. 1A**, green). CoF spots are identified as colocalized and co-moving nascent chain and folding spots.

**Fig. 1.**
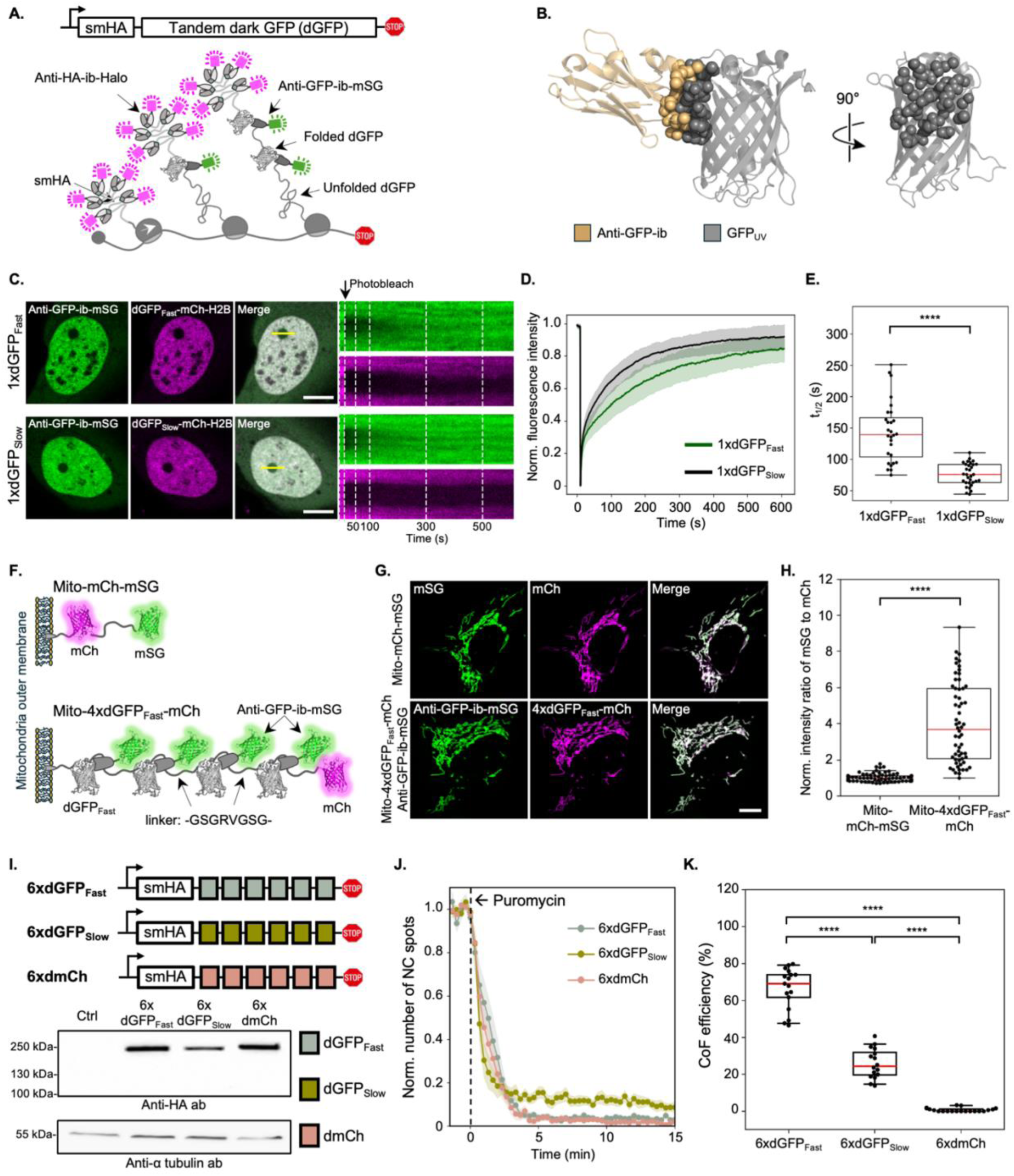
Tracking single-mRNA co-translational folding in live cells. **(A)** Schematic of the co-translational folding tracking (coTFT) technology. The coTFT reporter consists of an N-terminal smHA tag fused to tandem dark GFPs (dGFPs). Nascent chain (NC) spots are labeled by a co-expressed HaloTag-fused anti-HA intrabody (anti-HA-ib-Halo, magenta), stained with JF646 dyes, which binds the 10 HA epitopes within smHA. Correctly folded NCs are visualized by a conformation-sensitive anti-GFP intrabody fused to tandem mStayGold (anti-GFP-ib-mSG, green). Co-translational folding (coF) spots are identified by colocalization of NC (magenta) and folding (green) spots. **(B)** Crystal structure of anti-GFP-ib (LaG16, tan) bound to GFPuv (gray) (PDB: 6LR7), highlighting the interaction interface (spheres) and illustrating recognition of a conformational epitope. **(C)** Left, representative FRAP images showing the photobleached region of interest (ROI; 3 µm diameter dark circle) in nuclei of cells expressing anti-GFP-ib-mSG (green) and 1×dGFP-mCh-H2B (magenta). Right, kymographs displaying fluorescence recovery over time along transverse lines (yellow, 50 pixels) across the bleached ROI shown in the left panels. Top, 1×dGFP_Fast_; bottom, 1×dGFP_Slow_. **(D)** Normalized FRAP recovery curves of anti-GFP intrabody against dGFP_Fast_ and dGFP_Slow_ within the ROI. Green, dGFP_Fast_ (mean ± SEM, n = 30 cells); black, dGFP_Slow_ (mean ± SEM, n = 32 cells). **(E)** Box-and-whisker plots of the FRAP half recovery times. **(F)** Schematic of testing the designed linker for anti-GFP-ib-mSG binding to four tandem dGFP_Fast_ inserted between Mito and mCh (Mito-4×dGFP_Fast_-mCh). The control construct Mito-mCh-mSG is shown at the top. (**G)** Representative images of cells expressing Mito-mCh-mSG (top) or co-expressing Mito-4×dGFP_Fast_-mCh and anti-GFP-ib-mSG (bottom). (**H)** Box-and-whisker plots of normalized mitochondrial mSG to mCh fluorescence intensity ratios. From left to right: n = 74 cells (1.036 ± 0.027; mean ± SEM) and n = 60 cells (4.031 ± 0.287; mean ± SEM). (**I)** Schematic of the three coTFT reporters (6×dGFP_Fast_, 6×dGFP_Slow_, and 6×dmCh) and corresponding western blot analysis confirming full-length protein expression. α-Tubulin served as a loading control. **(J)** Puromycin assay of the three coTFT reporters (mean ± SEM; n = 10 cells for each reporter). (**K)** Box-and-whisker plots of coF efficiencies for 6×dGFP_Fast_ (66.55 ± 2.68%, n = 17 cells, 2,334 trajectories), 6×dGFP_Slow_ (25.84 ± 2.02%, n = 16 cells, 1,768 trajectories), and 6×dmCh (0.74 ± 0.24%, n = 20 cells, 2,987 trajectories) (mean ± SEM). In box-and-whisker plots, red lines indicate medians, **** p < 0.0001 (Mann-Whitney U test). Scale bars, 10 µm.

To determine whether coTFT can distinguish reporters with different folding kinetics, we selected fast- and slow-folding GFP variants: sfGFP and GFPuv (*20–22*) and quenched their fluorescence by introducing the G67A mutation (*18*, *23*), generating dGFP_Fast_ and dGFP_Slow_ reporters. We selected the high-affinity anti-GFP nanobody LaG16 (*24*) as the CS anti-GFP intrabody based on the crystal structure of the LaG16-GFPuv complex (PDB: 6LR7). The interaction interface spans multiple β-sheets on GFPuv (**Fig. 1B**), indicating that the epitope becomes accessible only after GFP adopts its folded conformation. Western blot analysis showed that the anti-GFP intrabody recognized the 6×dGFP_Fast_ reporter under native conditions but not after denaturation (**fig. S1A**), confirming its conformation-dependent binding and its ability to distinguish folding from non-folding nascent chains. Furthermore, the anti-GFP intrabody showed no detectable interaction with either smHA or mSG (**figs. S1B-C**), demonstrating its compatibility with the coTFT system.

To enable robust tracking of coF, the anti-GFP intrabody must bind both dGFP_Fast_ and dGFP_Slow_ with high affinity. We therefore characterized its binding kinetics in live cells using fluorescence recovery after photobleaching (FRAP) (*19*, *25*). In the FRAP assays, a defined nuclear region in cells co-expressing anti-GFP-ib-mSG and either 1×dGFP_Fast_- or 1×dGFP_Slow_-tagged mCh-H2B was photobleached and fluorescence recovery was monitored over time (**Fig. 1C**, left). Because mCh does not interact with the anti-GFP intrabody (**fig. S1B-C**), it serves as an appropriate fluorophore for the FRAP assays. Given the extremely slow turnover of H2B (*26*), fluorescence recovery in the mSG channel primarily reflects intrabody exchange kinetics, with slower recovery indicating tighter binding. As expected, the magenta channels (1×dGFP_Fast_- or 1×dGFP_Slow_-mCh-H2B) showed negligible recovery over ∼10 min, confirming the slow turnover of H2B (**Fig. 1C**, right). In contrast, the green channel (anti-GFP-ib-mSG) displayed different recovery kinetics for the dGFP variants (**Fig. 1D**). The half-recovery times (t**_1/2_**) of the anti-GFP intrabody were 143.92 ± 9.08 s for dGFP_Fast_ and 75.89 ± 3.18 s for dGFP_Slow_ (**Fig. 1E**), indicating slow turnover and therefore high-affinity binding to both variants. The slow turnover on dGFP_Slow_ is consistent with the reported high affinity (K_D_ = 6.7 nM) (*24*). The even slower recovery observed on dGFP_Fast_ suggests stronger binding, likely within the low-nanomolar or tighter range. Notably, the anti-GFP intrabody showed substantially slower recovery than the anti-HA intrabody (*27*), which is widely used for live-cell tracking of single-mRNA translation (*19*, *28–30*). Together, these results indicate that the anti-GFP intrabody binds both dGFP_Fast_ and dGFP_Slow_ with sufficient affinity to robustly label folding nascent chains as they emerge from the ribosome.

To increase fluorescence signals at coF sites, the reporter dGFP needs to be repeated multiple times using flexible linkers. We evaluated the linker design to determine whether it allowed maximal intrabody occupancy. In this assessment, cells co-expressing a mitochondrial outer membrane protein mitoNEET (Mito) (*31*) fused to 4×dGFP_Fast_-mCh (Mito-4×dGFP_Fast_-mCh) and anti-GFP-ib-mSG were imaged alongside control cells expressing Mito-mCh-mSG under identical conditions (**Fig. 1F**). Fluorescence intensities on mitochondria were quantified in both the mSG and mCh channels, and mSG/mCh ratio was normalized to the control (**Fig. 1G**). As shown in **Fig. 1H**, the normalized ratio increased from 1.036 ± 0.027 in control cells to 4.031 ± 0.287 in cells expressing Mito-4×dGFP_Fast_-mCh, indicating that each Mito-4×dGFP_Fast_-mCh molecule recruited approximately four anti-GFP intrabodies on average. These results demonstrate that the linker provides sufficient spacing to achieve near-maximal intrabody occupancy.

We next constructed three coTFT reporters: 6×dGFP_Fast_, 6×dGFP_Slow_, and 6×dark mCh (6×dmCh), using the validated linker, generating transcripts of identical length (**Fig. 1I**, top). Because the anti-GFP intrabody does not interact with mCh (**fig. S1B-C**), the 6×dmCh reporter serves as an appropriate negative control. All three reporters produced full-length proteins after 24 h of expression (**Fig. 1I**, bottom). Proper folding of the reporter proteins was verified using the corresponding bright reporters (6×bGFP_Fast_ and 6×bGFP_Slow_), in which fluorescence was restored by introducing the A67G mutation into each dGFP repeat (**figs. S1D-E**). Both bright reporters were excluded from the nucleus, consistent with their large size and supporting expression of intact full-length proteins. Aggregation was minimal for both reporters, although aggregates were observed somewhat more frequently in cells expressing 6×bGFP_Slow_. Importantly, co-expressed anti-GFP-ib-Halo did not colocalize with these aggregates (**fig. S1F**), indicating that the anti-GFP intrabody does not recognize aggregated reporter proteins.

Using the established coTFT technology, we successfully visualized live-cell coF spots from individual mRNAs in cells expressing 6×dGFP_Fast_ and 6×dGFP_Slow_, but not 6×dmCh. These spots were confirmed as bona fide coF spots by their rapid disappearance following puromycin treatment, which releases nascent chains from ribosomes (**Fig. 1J**). To ensure accurate quantification, only puromycin-responsive coF spots were included in all subsequent analyses. We defined coF efficiency as the fraction of nascent chain spots that colocalized with folding spots. The 6×dGFP_Fast_ reporter showed a significantly higher coF efficiency (66.55 ± 2.68%) than 6×dGFP_Slow_ (25.84 ± 2.02%), and both reporters displayed substantially higher coF efficiencies than the negative control 6×dmCh (0.74 ± 0.24%) (**Fig. 1K**). Together, these results establish coTFT as a platform for visualizing and quantifying coF on individual mRNAs in live cells and demonstrate its ability to distinguish proteins with distinct folding kinetics.

## Fast- and slow-folding reporters exhibit corresponding differences in translation elongation rates

We next applied coTFT to address a long-standing question in coF: how translation elongation and protein folding are coordinated. To investigate this relationship, we employed dGFP_Fast_ and dGFP_Slow_, two GFP variants with identical lengths and ∼97% protein sequence identity (**fig. S2A**) but distinct folding kinetics (*20–22*). To minimize confounding effects from codon usage and mRNA structure (*32*), we optimized their coding sequences to achieve ∼98% nucleotide sequence identity (**fig. S2B**). Each dGFP variant was repeated four times using the validated linker and followed by two tandem dmCh repeats, providing additional time for the last dGFP domain to fold and be detected by the anti-GFP intrabody before ribosome release. An identical N-terminal smHA tag enabled nascent chain tracking and 24× MS2 stem-loops in the 3’ untranslated region (3’UTR) allowed mRNA tethering when needed. This design (4×dGFP_Fast_ and 4×dGFP_Slow_, **Fig. 2A**) controls for transcript length, codon usage, mRNA structure, translation ramping, and overall sequence composition (∼99% nucleotide sequence identity and ∼98% protein sequence identity), allowing us to specifically interrogate the relationship between translation elongation and protein folding.

**Fig. 2.**
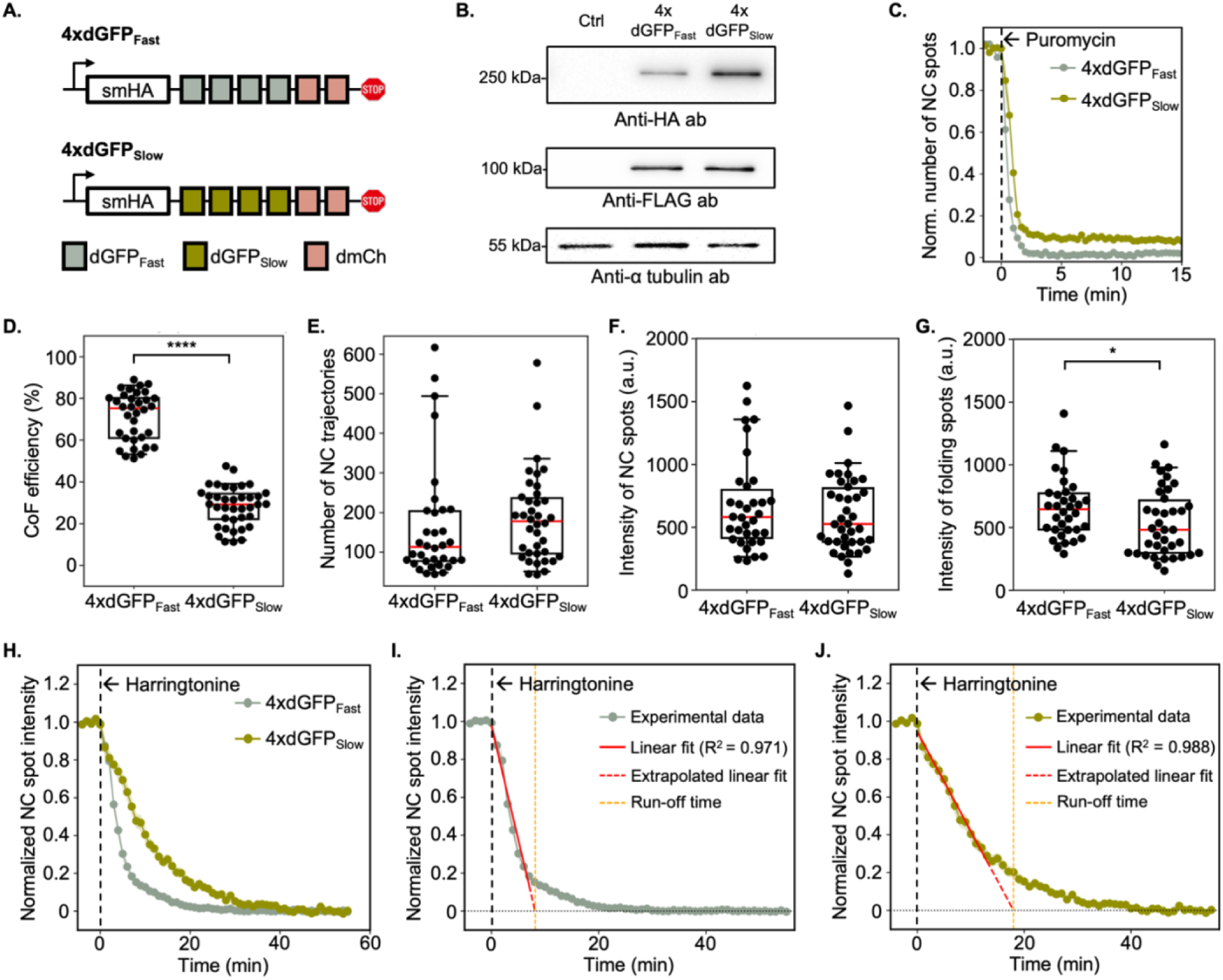
Fast- and slow-folding reporters exhibit corresponding differences in translation elongation rates. **(A)** Schematic of the 4×dGFP_Fast_ and 4×dGFP_Slow_ coTFT reporters, each consisting of an N-terminal smHA tag, four tandem dGFP_Fast_ or dGFP_Slow_ repeats, and two tandem dmCh repeats. **(B)** Western blot analysis of the 4×dGFP reporters expressed for 24 h, confirming full-length protein expression. 4×FLAG-mCh-β-actin served as a transfection control and α-tubulin as a loading control. **(C)** Puromycin assay results for the two coTFT reporters (mean ± SEM; n = 12 cells for 4×dGFP_Fast_ and n = 15 cells for 4×dGFP_Slow_). **(D)** Box-and-whisker plots of co-translational folding (coF) efficiencies for 4×dGFP_Fast_ (71.62 ± 2.06%, n = 34 cells, 5,598 trajectories) and 4×dGFP_Slow_ (28.00 ± 1.49%, n = 38 cells, 7,077 trajectories). **(E)** Box-and-whisker plots of the number of nascent chain (NC) trajectories per cell corresponding to the data in (D). **(F)** Box-and-whisker plots of the median NC intensity per cell corresponding to the data in (D). **(G)** Box-and-whisker plots of the median folding spot intensity per cell for 4×dGFP_Fast_ (4,259 trajectories) and 4×dGFP_Slow_ (2,097 trajectories) corresponding to the data in (D). **(H)** Harringtonine run-off assay results for 4×dGFP_Fast_ (n = 26 cells) and 4×dGFP_Slow_ (n = 28 cells). **(I)** Linear fit of the harringtonine run-off curve for 4×dGFP_Fast_, resulting in a run-off time of 8.1 min and an elongation rate of 3.878 aa/s. **(J)** Linear fit of the harringtonine run-off curve for 4×dGFP_Slow_, resulting in a run-off time of 18.0 min and an elongation rate of 1.629 aa/s. In box-and-whisker plots, red lines indicate medians. * p < 0.05, **** p < 0.0001 (Mann-Whitney U test).

We confirmed by western blot that both reporters produced full-length proteins (**Fig. 2B**). Proper folding was verified using the corresponding bright reporters (4×bGFP_Fast_ and 4×bGFP_Slow_), which exhibited robust green fluorescence in live cells (**figs. S3A-B**). Similar to the 6×bGFP reporters, both bright 4×bGFP reporters were excluded from the nucleus, consistent with their large size and supporting the integrity of the full-length protein (**figs. S3A-B**).

For coF imaging, we generated a monoclonal stable cell line co-expressing anti-HA-ib-Halo and anti-GFP-ib-mSG under the control of a tetracycline-inducible promoter to minimize cell-to-cell variation in intrabody expression. Both reporters produced puromycin responsive coF spots (**Fig. 2C)**. Consistent with the results obtained using 6×dGFP, the two reporters exhibited distinct coF efficiencies (**movies S1** and **S2**), with 4×dGFP_Fast_ displaying a substantially higher coF efficiency (71.62 ± 2.06%) than 4×dGFP_Slow_ (28.00 ± 1.49%) (**Fig. 2D**). The two reporters exhibited comparable numbers and intensities of nascent chains (**Figs. 2E-F**), whereas 4×dGFP_Fast_ displayed modestly brighter folding spots (**Fig. 2G**). This difference may arise from more folded dGFP domains on translating ribosomes in the 4×dGFP_Fast_ reporter, which was confirmed later in this study.

Harringtonine run-off assays revealed that the 4×dGFP_Fast_ reporter exhibited a more rapid decay than 4×dGFP_Slow_, indicating faster translation elongation (**Fig. 2H**). Linear fitting yielded elongation rates of 3.878 aa/s for 4×dGFP_Fast_ and 1.629 aa/s for 4×dGFP_Slow_ (**Figs. 2I-J**). Because other major determinants of elongation were largely controlled in our reporter design and the presence of the anti-GFP intrabody did not affect elongation (**figs. S3C-D**), the observed elongation difference likely reflects the distinct folding kinetics of the reporters, suggesting that protein folding can influence translation elongation.

## Perturbing translation elongation alters co-translational folding outcomes

Having established that protein folding can influence translation elongation during coF, we next investigated whether altering elongation affects folding outcomes. To perturb elongation, we employed the translation-pausing peptide XBP1u (S255A mutant, 26 amino acids) (*33–36*), which inhibits peptide-bond formation and thereby slows translation elongation. XBP1u was inserted either immediately upstream of the stop codon (4×dGFP_Fast_-XBP1u) or between the N-terminal smHA tag and the first dGFP_Fast_ domain (XBP1u-4×dGFP_Fast_) within the 4×dGFP_Fast_ reporter (**Fig. 3A**).

**Fig. 3.**
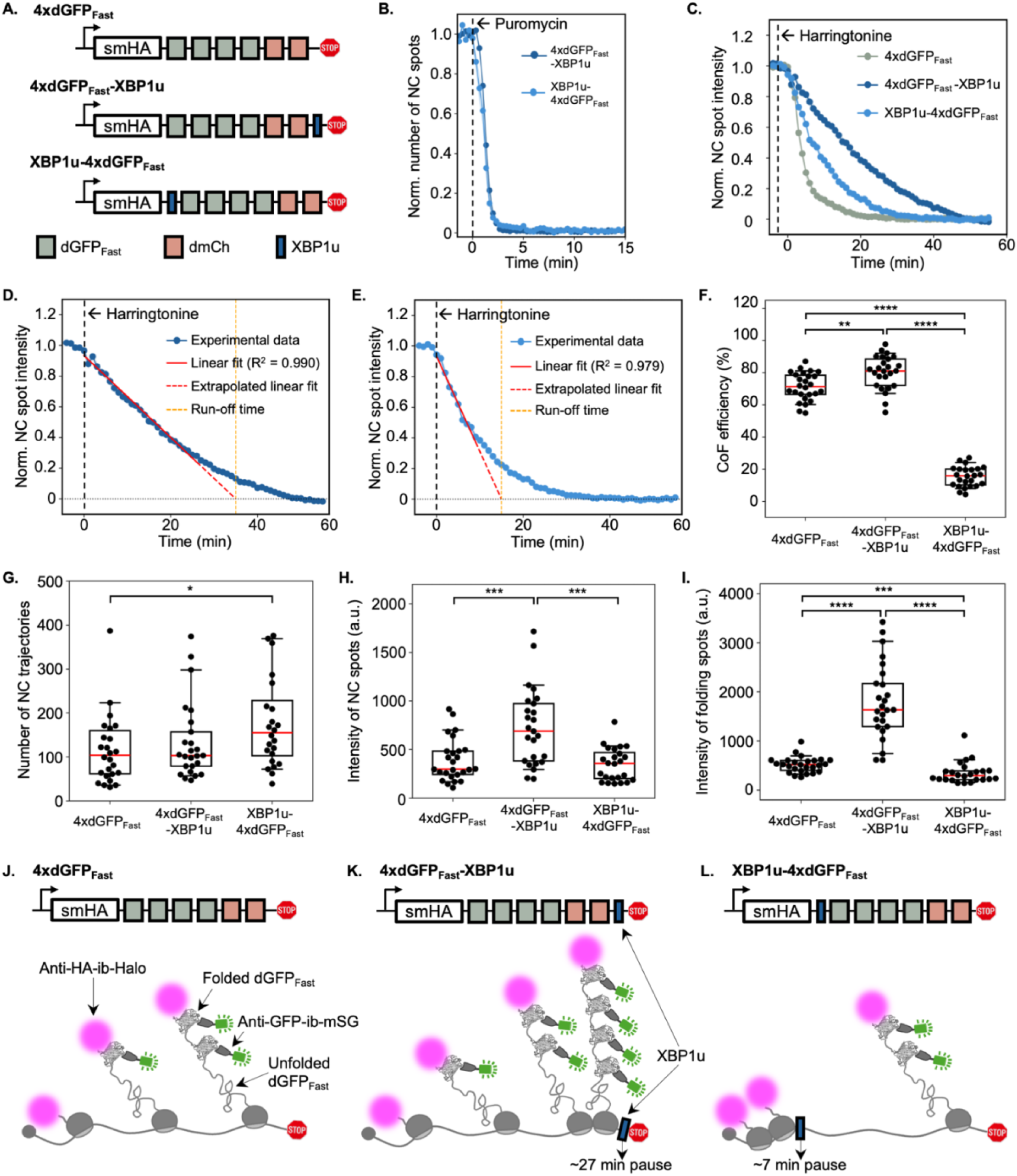
Perturbing translation elongation alters co-translational folding outcomes. **(A)** Schematic of the 4×dGFP_Fast_ reporters with or without the XBP1u translation pausing peptide. **(B)** Puromycin assay results for 4×dGFP_Fast_-XBP1u (n = 12 cells) and XBP1u-4×dGFP_Fast_ (n = 13 cells). **(C)** Harringtonine run-off assay results for 4×dGFP_Fast_ (n = 26 cells, replotted from Figs. 2H and 2I), 4×dGFP_Fast_-XBP1u (n = 22 cells), and XBP1u-4×dGFP_Fast_ (n = 24 cells). **(D)** Linear fit of the harringtonine run-off curve for 4×dGFP_Fast_-XBP1u, resulting in a run-off time of 35.0 min and an elongation rate of 0.826 aa/s. **(E)** Linear fit of the harringtonine run-off curve for XBP1u-4×dGFP_Fast_, resulting in a run-off time of 15.0 min and an elongation rate of 2.012 aa/s. **(F)** Box-and-whisker plots of co-translational folding (coF) efficiencies for 4×dGFP_Fast_ (71.38 ± 1.69%, n = 26 cells, 3,357 trajectories), 4×dGFP_Fast_-XBP1u (80.28 ± 2.06%, n = 25 cells, 3,343 trajectories), and XBP1u-4×dGFP_Fast_ (15.25 ± 1.26%, n = 24 cells, 4,457 trajectories). **(G)** Box-and-whisker plots of the number of nascent chain (NC) trajectories per cell corresponding to the data in (F). **(H)** Box-and-whisker plots of the median NC intensity per cell corresponding to the data in (F). **(I)** Box-and-whisker plots of the median folding spot intensity per cell for 4×dGFP_Fast_ (2,475 trajectories), 4×dGFP_Fast_-XBP1u (2,733 trajectories), and XBP1u-4×dGFP_Fast_ (675 trajectories) corresponding to the data in (F). **(J-L)** Models illustrating the expected ribosome density and spatial distribution on XBP1u-containing reporters relative to the parental 4×dGFP_Fast_ reporter. In box-and-whisker plots, red lines indicate medians, * p < 0.05, ** p < 0.01, *** p < 0.001, **** p < 0.0001 (Mann-Whitney U test).

We first confirmed that reporters containing XBP1u at either position produced full-length proteins, although at substantially lower levels than the reporter lacking XBP1u (**fig. S4A**). Puromycin treatment validated that the observed spots in XBP1u reporters represented coF events, as they rapidly disappeared upon puromycin treatment (**Fig. 3B**). To verify that XBP1u insertion did not impair GFP folding, we generated the corresponding bright reporters, 4×bGFP_Fast_-XBP1u and XBP1u-4×bGFP_Fast_, and successfully visualized green fluorescent cells expressing either reporter (**figs. S4B-C**). Notably, we observed puromycin-responsive nascent chain spots illuminated by folded and fluorescent GFP in cells expressing 4×bGFP_Fast_-XBP1u. Given that bGFP_Fast_ requires ∼10 min to fold and fluoresce, this observation suggests that XBP1u induces a substantial translational pause, allowing fluorescent GFP to accumulate while the nascent chain remains tethered to the mRNA.

We next measured the translation elongation rates of XBP1u-4×dGFP_Fast_ and 4×dGFP_Fast_-XBP1u using harringtonine run-off assays. As shown in **Fig. 3C**, XBP1u-4×dGFP_Fast_ displayed a run-off time ∼6.9 min longer than that of 4×dGFP_Fast_ (15.0 min versus 8.1 min), consistent with the ∼6 min pause previously measured by socRNA (*36*). Interestingly, 4×dGFP_Fast_-XBP1u showed an even longer run-off time than XBP1u-4×dGFP_Fast_ (35.0 min versus 15.0 min). The additional ∼20 min delay may arise from the clearance of a ribosome traffic jam near the stop codon. Linear fitting of the run-off curves yielded elongation rates of 0.826 aa/s for 4×dGFP_Fast_-XBP1u (**Fig. 3D**) and 2.012 aa/s for XBP1u-4×dGFP_Fast_ (**Fig. 3E**), both significantly slower than that of 4×dGFP_Fast_ (3.878 aa/s). These results demonstrate that both the presence and position of XBP1u strongly influence translation elongation.

Then, we quantified coF efficiencies for these reporters. As expected, 4×dGFP_Fast_-XBP1u exhibited a significantly higher coF efficiency than 4×dGFP_Fast_, increasing from 71.38 ± 1.69% to 80.28 ± 2.06% (**Fig. 3F**). This increase is most likely due to prolonged ribosome occupancy on the mRNA, which extends the observation window for coF events. In contrast, XBP1u-4×dGFP_Fast_ exhibited a markedly reduced coF efficiency (15.25 ± 1.26%) (**Fig. 3F**). Although similar numbers of nascent chains were detected for 4×dGFP_Fast_-XBP1u compared to 4×dGFP_Fast_ and XBP1u-4×dGFP_Fast_ (**Fig. 3G**), both nascent chain and folding spot intensities were significantly higher for 4×dGFP_Fast_-XBP1u (**Figs. 3H-I**). The elevated intensities are consistent with delayed ribosome release and increased ribosome occupancy relative to the other two reporters (**Figs. 3J-K**). In contrast, XBP1u-4×dGFP_Fast_ exhibited modestly more nascent chains than 4×dGFP_Fast_ (**Fig. 3G**), while nascent chain intensities remained comparable (**Fig. 3H**), suggesting similar ribosome loading (**Figs. 3J, 3L**). However, the substantially dimmer folding spots in XBP1u-4×dGFP_Fast_ (**Fig. 3I**) indicate a marked reduction in coF events per mRNA, likely due to a lower density of translating ribosomes downstream of the XBP1u sequence (**Figs. 3J, 3L**).

We applied the same XBP1u strategy to the 4×dGFP_Slow_ reporter (**fig. S4D**). Consistent with the results obtained for the 4×dGFP_Fast_ reporters, XBP1u-containing reporters produced full-length proteins but at reduced abundance (**fig. S4E**). Quantification of puromycin-validated coF spots (**fig. S4F**) revealed that 4×dGFP_Slow_-XBP1u exhibited a markedly higher coF efficiency (70.04 ± 2.61%), whereas XBP1u-4×dGFP_Slow_ displayed a substantially lower coF efficiency (7.47 ± 0.71%), than the parental 4×dGFP_Slow_ reporter (29.67 ± 1.61%) (**fig. S4G**). Across all three reporters, nascent chain and folding spot intensities displayed trends comparable to those observed for the 4×dGFP_Fast_ reporters (**figs. S4H-J**).

Taken together, these results demonstrate that perturbing translation elongation substantially reshapes coF outcomes by altering the frequency of coF events on individual mRNAs. They further establish coTFT as a live-cell platform for real-time analysis of coF dynamics.

## CoTFT reporters exhibit distinct co-translational folding kinetics in live cells

In developing coTFT, we selected dGFP_Fast_ and dGFP_Slow_ based on their distinct *in vitro* refolding kinetics (*20–22*). However, their folding kinetics on translating ribosomes in live cells had not been quantitatively characterized. To address this knowledge gap, we generated a series of reporters containing increasing numbers (1× ∼ 6×) of dGFP_Fast_ (**Fig. 4A**) or dGFP_Slow_ (**Fig. 4B**) repeats and assessed how many dGFP domains could fold before translation termination.

**Fig. 4.**
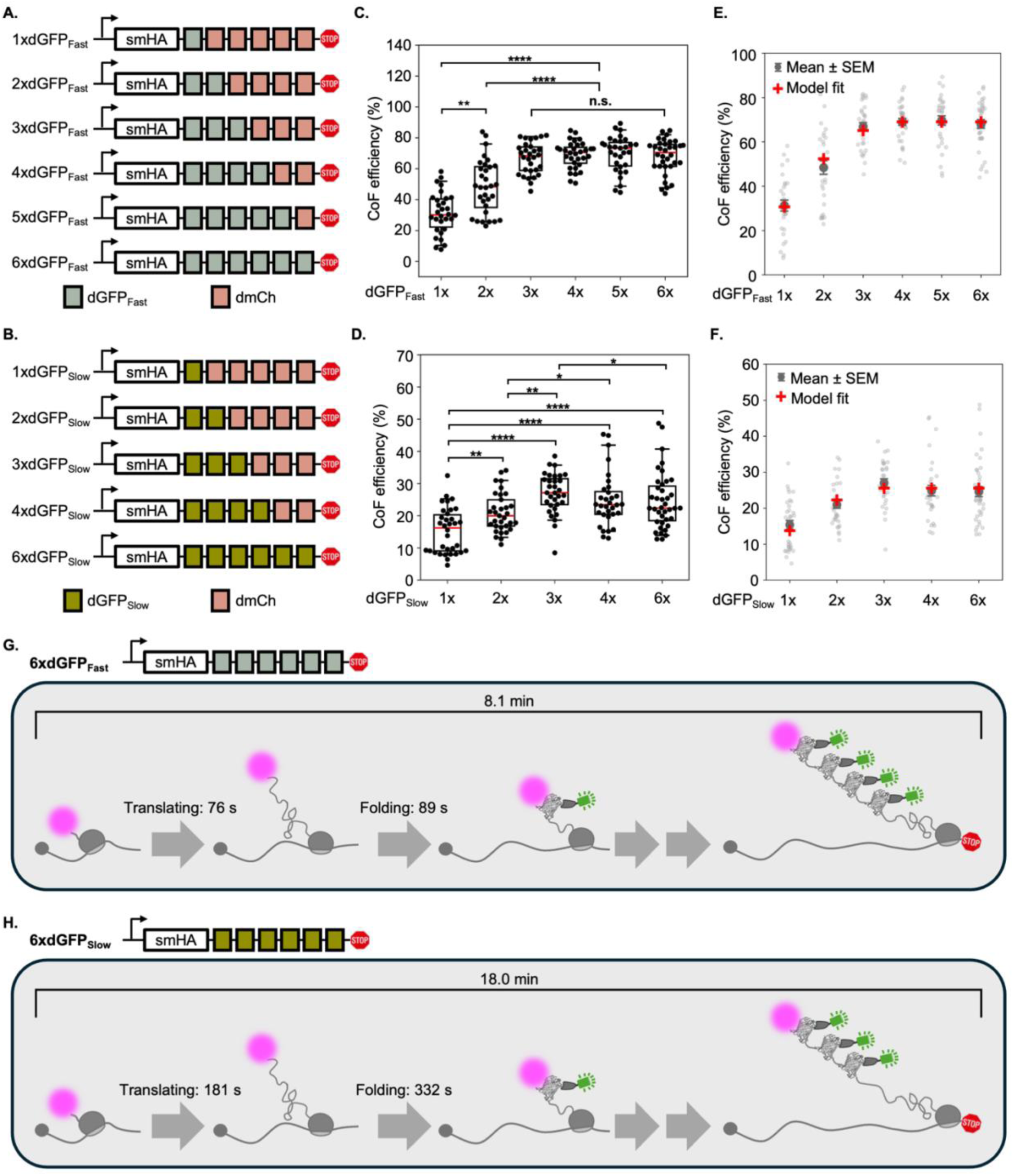
CoTFT reporters exhibit distinct co-translational folding kinetics in live cells. **(A-B)** Schematic of the dGFP_Fast_ (A) and dGFP_Slow_ (B) reporter series containing increasing numbers of dGFP repeats (1× - 6×). **(C-D)** Box-and-whisker plots showing co-translational folding (coF) efficiencies of the dGFP_Fast_ reporter series in (C) (1×dGFP_Fast_: 31.29 ± 2.44%, n = 30 cells, 3,611 trajectories; 2×dGFP_Fast_: 48.43 ± 3.06%, n = 32 cells, 3,702 trajectories; 3×dGFP_Fast_: 66.60 ± 1.87%, n = 30 cells, 4,767 trajectories; 4×dGFP_Fast_: 69.04 ± 1.52%, n = 32 cells, 3,777 trajectories; 5×dGFP_Fast_: 69.65 ± 2.07%, n = 31 cells, 2,948 trajectories; 6×dGFP_Fast_: 68.07 ± 1.91%, n = 34 cells, 4,331 trajectories) and the dGFP_Slow_ reporter series in (D) (1×dGFP_Slow_: 15.36 ± 1.20%, n = 34 cells, 4,171 trajectories; 2×dGFP_Slow_: 21.02 ± 1.06%, n = 33 cells, 4,414 trajectories; 3×dGFP_Slow_: 26.98 ± 1.11%, n = 32 cells, 3,825 trajectories; 4×dGFP_Slow_: 24.86 ± 1.47%, n = 32 cells, 4,533 trajectories; 6×dGFP_Slow_: 24.61 ± 1.40%, n = 39 cells, 4,489 trajectories). **(E-F)** Fits of the folding time model to the experimentally measured coF efficiencies of the dGFP_Fast_ (E) and dGFP_Slow_ (F) reporter series. **(G)** Predicted folding time of the 6×dGFP_Fast_ reporter, indicating that up to four dGFP_Fast_ repeats can fold before translation termination. **(H)** Predicted folding time of the 6×dGFP_Slow_ reporter, indicating up to three dGFP_Slow_ repeats can fold before translation termination. In box-and-whisker plots, red lines indicate medians. * p < 0.05, ** p < 0.01, **** p < 0.0001 (Mann-Whitney U test).

We first confirmed by western blot that all reporters produced full-length proteins (**figs. S5A-B**). We then quantified coF efficiency for each reporter (**Figs. 4C-D**). For both dGFP_Fast_ and dGFP_Slow_ reporter series, coF efficiency increased as the number of dGFP repeats increased from 1× to 3×, followed by a plateau at higher repeat numbers (3× ∼ 6×) (modestly decline between 3× and 6×dGFP_Slow_, p < 0.05). These results suggest that only the first three to four dGFP repeats have sufficient time to fold and be recognized by the anti-GFP intrabody before translation termination.

Fitting a mathematical model to the experimentally measured coF efficiencies (**Figs. 4E-F**) yielded estimated folding times of approximately 89 s for dGFP_Fast_ and 332 s for dGFP_Slow_ (see Materials and Methods section), demonstrating that dGFP_Fast_ folds substantially faster than dGFP_Slow_ on translating ribosomes in live cells. Using this model, we predict that the first four dGFP_Fast_ domains can fold before translation termination (**Fig. 4G** and **table S2**). However, the fourth folded dGFP_Fast_ domain is predicted to remain detectable for only a short time window (∼27 s) before ribosome release, resulting in little contribution to the coF efficiency measurements and explaining the absence of a significant increase in coF efficiency between the 3× and 4×dGFP_Fast_ reporters. In contrast, the model predicts that only the first three dGFP_Slow_ domains can fold before translation termination (**Fig. 4H and table S2**). This prediction is consistent with the dimmer folding spots observed for 4×dGFP_Slow_ relative to 4×dGFP_Fast_ (**Fig. 2G**).

Together, these results demonstrate that coTFT can quantitatively resolve protein folding time during coF in live cells. The estimated folding times confirm that dGFP_Fast_ and dGFP_Slow_ exhibit distinct folding kinetics on translating ribosomes in live cells, consistent with their previously reported relative folding behaviors *in vitro* (*20–22*).

## Discussion

CoF couples protein synthesis with the acquisition of native protein structure, yet direct observation of folding on translating ribosomes in live cells has remained elusive. By developing coTFT, we provide an imaging platform that enables visualization and quantification of coF at the level of individual mRNAs in live cells. This capability allows mechanistic investigation of how translation elongation and folding are coordinated within their native cellular context.

Using coTFT, we found that fast- and slow-folding reporters, which differed by 3.7-fold in folding times, exhibited a corresponding 2.4-fold difference in translation elongation rates despite possessing nearly identical nucleotide and protein sequences and identical transcript lengths. The ∼1.8% difference in protein sequence is unlikely to account for the observed elongation difference, as formation of the peptide bonds involving the substituted amino acids in dGFP_Fast_ and dGFP_Slow_ (**fig. S2A**) is generally much faster than the overall elongation cycle and therefore is not rate-limiting (*32*, *37*). Likewise, the nucleotide sequences differ by only 0.8% and contain highly optimized codons (CAI = 0.91, **fig. S2B**), indicating that codon usage is not a major contributor to the difference in translation elongation. Although subtle contributions from codon usage, mRNA structure, or sequence composition cannot be excluded, our findings suggest that protein folding can, in turn, influence elongation.

The mechanism underlying this feedback, however, is unlikely to involve folding-generated pulling forces near the ribosome exit tunnel (*13*), as kinetic analyses indicate that both reporters fold well beyond the tunnel exit (∼345 aa for dGFP_Fast_ and ∼542 aa for dGFP_Slow_). Instead, the distinct elongation rates may reflect differential interactions between folding nascent chains and the ribosome. Ribosomes are increasingly recognized as active participants in coF, interacting with nascent chains to stabilize or destabilize folding intermediates (*12*, *38–42*). Such interactions may be more prolonged or frequent for the slow-folding reporter, thereby slowing elongation. Alternatively, the two reporters may differentially engage ribosome-associated chaperones, such as the nascent polypeptide-associated complex (NAC) (*43*, *44*). By interacting with emerging nascent chains near the ribosome exit tunnel, NAC may modulate folding intermediates and thereby influence elongation. The molecular basis of this feedback remains to be elucidated, and the coTFT technology developed in this study should enable dissection of these mechanisms.

We further showed that altering translation elongation substantially changed coF outcomes, providing direct evidence that elongation influences folding. Notably, XBP1u-containing reporters produced markedly lower amounts of full-length protein despite exhibiting either increased or decreased coF efficiencies. Because prolonged XBP1u-mediated pausing can trigger ribosome collisions and activate ribosome-associated quality-control pathways (*45*, *46*), the reduction in protein production is more likely attributable to collision-induced translational surveillance than to altered coF efficiencies.

Together, our work not only establishes coTFT as a novel technology for studying coF with single-mRNA resolution in live cells, but also provides direct evidence for a new regulatory principle in which translation elongation and protein folding are bidirectionally coupled to coordinate co-translational protein biogenesis in live cells.

## Acknowledgments

We thank all members of the Zhao lab for valuable feedback on the manuscript. We also thank Drs. Timothy Stasevich, Judith Frydman, Hideki Taguchi, and Olivia Rissland for insightful discussion. This study was supported in part by the National Institutes of Health P30CA046934 funded Cancer Center Flow Cytometry Shared Resource [RRID:SCR_022035].

## Funding

This work was supported by NIH grants K99GM141453 (NZ), R00GM141453 (NZ), R35GM160021(NZ), and Cystic Fibrosis Foundation grant 005749A123 (NZ).

## Author contributions

Conceptualization: RMS, LUA, NZ

Methodology: RMS, LUA, NZ

Investigation: RMS, LUA, TMB, NZ

Visualization: RMS, LUA, NZ

Funding acquisition: NZ

Project administration: NZ

Supervision: NZ

Writing – original draft: RMS, LUA, NZ

Writing – review & editing: RMS, LUA, TMB, NZ

## Competing interests

Authors declare that they have no competing interests.

## Data, code, and materials availability

The complete code used for this study is available in GitHub: https://github.com/ningzhaoAnschutz/cof_paper.git.

Microscopy images available at: 10.5281/zenodo.20936426.

## Notes

### Competing Interest Statement

The authors have declared no competing interest.

