## Supplemental Information for "Live-cell co-translational folding tracking reveals bidirectional coupling between translation and folding"

**The PDF file includes:**

Materials and Methods
Figs. S1 to S6
Table S2

**Other Supplementary Materials for this manuscript include the following:**

Table S1
Movies S1 to S2

**Movie S1. Co-translational folding tracking of the 4×dGFP$_{Fast}$ reporter.** Representative live-cell imaging of a cell co-expressing the 4×dGFP$_{Fast}$ reporter with the cognate imaging intrabodies, showing nascent chain spots (magenta) and folding spots (green). Colocalized, comoving magenta and green spots represent co-translational folding spots. The movie is related to Figs 2D-G.

**Movie S2. Co-translational folding tracking of the 4×dGFP$_{Slow}$ reporter.** Representative live-cell imaging of a cell co-expressing the 4×dGFP$_{Slow}$ reporter with the cognate imaging intrabodies, showing nascent chain spots (magenta) and folding spots (green). Colocalized, comoving magenta and green spots represent co-translational folding spots. The movie is related to Figs 2D-G.

### Materials and Methods

#### *Plasmids*

All plasmids used in this study are listed in table S1. Cloning was designed in SnapGene (version 8.0.1 or later) using synthetic DNA fragments from Twist Bioscience and primers from Invitrogen. PCR amplification was performed using the Expand High Fidelity PCR System (Roche), and restriction enzymes were obtained from New England Biolabs. The PiggyBac-Tet-On vector was generated by VectorBuilder. All constructs were verified by whole-plasmid nanopore sequencing (Quintara Biosciences).

#### *Cell culture*

Human U-2 OS cells (ATCC HTB-96) and house-made U-2 OS stable cell lines were cultured in DMEM (Gibco, 11960-044) supplemented with 10% fetal bovine serum (Avantor, 97068-085, lot 128K24), 2 mM L-glutamine (Gibco, 25030081), and 1% penicillin-streptomycin (Gibco, 15140122). Cells were maintained at 37 °C in a humidified incubator with 5% $CO_2$.

#### *Transfection*

Plasmids were prepared using the ZymoPURE II Plasmid Midiprep Kit (Zymo Research) and diluted in ultrapure water to approximately 100 ng/µL. Prior to transfection, U-2 OS cells seeded in 35-mm MatTek imaging dishes or standard tissue culture dishes were washed twice with PBS (pH 7.4) and incubated in prewarmed Opti-MEM (Gibco) supplemented with 10% fetal bovine serum (Avantor). Cells were transfected with 1.25 or 2.5 µg plasmid DNA using Lipofectamine LTX and PLUS reagents (Thermo Fisher Scientific) according to the manufacturer's instructions. After 3 h, the transfection medium was replaced with DMEM (Gibco) supplemented with 10% fetal bovine serum (Avantor), 2 mM L-glutamine (Gibco), and 1% penicillin-streptomycin (Gibco). Cells were maintained at 37 °C in a humidified atmosphere containing 5% $CO_2$ until imaging or harvesting.

#### *Live-cell imaging conditions*

Live-cell imaging was performed on a Leica Stellaris 5 confocal microscope using the acquisition settings described previously (*27*). Briefly, cells were maintained at 37 °C with 5% $CO_2$ and 51% relative humidity during imaging. Images were acquired with a 63× oil-immersion objective at 16-bit depth with a field of view of 512 × 512 pixels (129.89 nm per pixel). For z-stack acquisition, images were collected at 0.3-µm steps. Image acquisition was controlled using Leica Application Suite X (LAS X, version 4.5.0.25531).

Unless otherwise indicated, live-cell imaging was performed in resonance scanning mode (8 kHz) with bidirectional scanning, line averaging of 16, a bidirectional phase offset of −87.74, and a zoom factor of 2.78. For co-translational folding efficiency measurements, z-stack movies were acquired every 5 s for 20 frames using a 488-nm laser at 2% power (emission, 500–550 nm) and a 638-nm laser at 5% power (emission, 645–700 nm). For puromycin assays, imaging was performed under the same settings in multi-position acquisition mode every 20 s for 60 frames, enabling simultaneous imaging of multiple cells per chamber. Puromycin (100 µg/mL; Goldbio) was added immediately after acquisition of the fifth frame. For harringtonine run-off assays, only the 638-nm laser was used (5% power; emission, 645–700 nm). Z-stack images were acquired every 1 min for 60 frames in multi-position mode, and harringtonine (3 µg/mL; Cayman Chemical) was added immediately after acquisition of the fifth frame.

FRAP experiments were performed using single-plane time-lapse imaging in FRAP mode with a scan speed of 800 Hz, a zoom factor of 2.45, and line averaging of 1. Fluorescence signals were detected using a HyD S detector operating in photon-counting mode. Samples were excited with 488-nm and 561-nm lasers at 2% power. Photobleaching was performed using the 488-nm laser at 30% power in Zoom-in bleaching mode. The bleached region of interest (ROI) was defined as a circular area with a diameter of 3 µm. Each FRAP acquisition consisted of 10 pre-bleach frames collected at 1-s intervals, followed by 300 post-bleach frames acquired at 2-s intervals.

***Fluorescence recovery after photobleaching***

For FRAP imaging, U-2 OS cells were seeded in MatTek chambers and transiently co-transfected with 1.25 µg of anti-GFP-ib-mSG (table S1) and 1.25 µg of either 1×dGFP$_{Fast}$-mCh-H2B (table S1) or 1×dGFP$_{Slow}$-mCh-H2B (table S1). FRAP experiments were performed 20 h post-transfection.

FRAP data were analyzed using a custom Python pipeline implemented in MicroLive (*47*). The pipeline reads Leica .lif files, extracts image stacks, time intervals, and pixel size metadata, and concatenates pre-photobleaching (Pre) and post-photobleaching (Pb1; 0.5 fps) acquisitions into a single time-lapse movie. Cell nuclei were segmented and FRAP ROIs were automatically identified by detecting the bleached spot between pre- and post-bleach frames or through threshold-based intensity tracking. Fluorescence intensities within the ROI were background-subtracted and quantified over time to generate recovery curves. Recovery curves that approached pre-bleach fluorescence levels were fitted with a single-exponential model to estimate fluorescence recovery half-times ($t_{1/2}$). ROIs with poor detection or segmentation were excluded after manual inspection.

***Stable cell lines generation***

To enable robust tracking of co-translational folding in live cells, we generated a monoclonal U-2 OS cell line expressing anti-HA-ib-Halo and anti-GFP-ib-mSG under the control of a Tet-On inducible promoter. U-2 OS cells were transiently co-transfected with PiggyBac-Tet-On-intrabodies (**table S1**) plasmid and a Super PiggyBac Transposase (NovoPro, V012800) at a 4:1 ratio. The culture medium was replaced 6 h after transfection. Hygromycin selection (350 µg/mL) was initiated 48 h post transfection and maintained for 14 days. Survived cells were subsequently subjected to single-cell fluorescence-activated cell sorting into 96-well plates to generate monoclonal cell lines. Individual clones were expanded and screened for inducible intrabody expression and performance in co-translational folding tracking. A single clone that exhibited robust and reproducible co-translational folding tracking was selected for all co-translational folding tracking imaging in this study.

For harringtonine run-off assays, we generated a stable cell line for tracking membrane-tethered co-translational folding spots through the interaction between the 24× MS2 stem-loops in the 3'UTR of each coTFT reporter and a cognate tandem MS2 coat protein fused to a CAAX membrane-targeting domain (tdMCP-CAAX). The monoclonal stable cell line described above was transduced with third-generation lentivirus encoding tdMCP-CAAX using the Lenti-X™ Packaging Single Shots system (Takara). To produce lentivirus, HEK293T cells (a generous gift from the laboratory of Joe Nassour) were transfected with the Lenti-tdMCP-CAAX plasmid (**table S1**) using Lenti-X™ Packaging Single Shots (VSV-G; Takara). The culture medium was replaced 6 h after transfection. Viral supernatants were collected 24 h later, passed through a 0.45 µm filter, and concentrated overnight at 4 °C using Lenti-X™ Concentrator (Takara). The above established monoclonal stable cells were incubated with concentrated lentivirus supplemented with 2 µg/mL polybrene (Sigma-Aldrich) for 24 h. Transduced cells were subsequently selected with blasticidin (10 µg/mL) for 14 days. Individual clones were isolated, expanded, and screened for efficient reporter tethering. A single clone exhibiting robust and reproducible reporter tethering was selected for all harringtonine run-off assays in this study.

***Western blotting***

For analysis of coTFT reporter expression, cells seeded in 35 mm culture dishes at 75% confluency were transiently co-transfected with 23 µL of the indicated coTFT reporter plasmid (100 ng/µL) and 2 µL of a 4×FLAG-mCh-β-actin (**table S1**) plasmid (100 ng/µL) as a transfection control. Cells were harvested 24 h post-transfection and lysed in 150 µL Pierce™ IP Lysis Buffer (Thermo Fisher Scientific) supplemented with cOmplete™ Protease Inhibitor (Roche). Total protein concentrations were determined using the Pierce™ BCA Protein Assay Kit (Thermo Fisher Scientific). For western blot analysis, 20 µg of total protein from each lysate was mixed with sample buffer (Bio-Rad, 161-0737), incubated at 95 °C for 15 min, and loaded onto a 4-20% Mini-PROTEAN® TGX™ Precast Gel (Bio-Rad). Gels were run in an SDS running buffer (25 mM Tris, 192 mM glycine, 0.1% SDS) at 105 V for 5 min, followed by 200 V for at least 40 min to achieve protein separation. Proteins were subsequently transferred onto a 0.22 µm nitrocellulose membrane (Cytiva) at 105 V for 35 min using wet transfer with Towbin buffer (25 mM Tris, 192 mM glycine, 20% methanol). Membranes were blocked for 1 h at room temperature in a blocking buffer consisting of 5% non-fat milk powder in TBST (0.05% Tween-20 (Thermo Fisher Scientific) in TBS, pH 7.4). To detect coTFT reporters, membranes were incubated overnight at room temperature with mouse monoclonal anti-HA antibody (12CA5; Roche, ROAHA) diluted 1:5000 in blocking buffer, followed by incubation with HRP-conjugated anti-mouse secondary antibody (Cell Signaling Technology, 7076S; 1:5000 dilution in blocking buffer) for 1 h at room temperature. As a loading control, α-tubulin was detected sequentially using rabbit polyclonal anti-α-tubulin antibody (Cell Signaling Technology, 2144S; 1:5000 dilution in blocking buffer) incubated overnight at room temperature, followed by HRP-conjugated anti-rabbit secondary antibody (Cell Signaling Technology, 7074S; 1:5000 dilution in blocking buffer) for 1 h at room temperature. Lastly, to detect the transfection control, membranes were subsequently incubated overnight at room temperature with mouse monoclonal anti-FLAG antibody (9A3; Cell Signaling Technology, 8146S; 1:5000 dilution in blocking buffer), followed by HRP-conjugated anti-mouse secondary antibody (Cell Signaling Technology, 7076S; 1:5000 dilution in blocking buffer) for 1 h at room temperature. Blots were developed using Clarity™ Western ECL Substrate (Bio-Rad) and imaged on a ChemiDoc MP Imaging System (Bio-Rad).

For native western blot analysis, cells were transiently transfected with 25 µL of the indicated coTFT reporter plasmid (100 ng/µL) and harvested 12 h post-transfection. To preserve native protein conformations, 10 µg of total cell lysate was mixed with Native Sample Buffer (Bio-Rad, 161-0738) and loaded onto a 4-20% Mini-PROTEAN® TGX™ Precast Gel (Bio-Rad). Gels were run in native running buffer (25 mM Tris, 192 mM glycine) at 4 °C, initially at 105 V for 5 min, followed by 150 V for at least 60 min to achieve protein separation. Proteins were subsequently transferred onto a 0.22 µm nitrocellulose membrane (Cytiva) at 105 V for 35 min at 4 °C using wet transfer with Towbin buffer. To detect native folded GFP reporters, the membrane was incubated overnight at 4 °C in diluted M13 phage particles displaying the anti-GFP intrabody in blocking buffer (50 uL of phage particles with a titer of 4.8 x $10^{13}$ cfu/mL). The membrane was subsequently incubated for 1 h at 4 °C with HRP-conjugated anti-M13 bacteriophage secondary antibody (Sino Biological, 11973-MM05T-H; 1:5000 dilution in blocking buffer). Blots were developed using Clarity™ Western ECL Substrate (Bio-Rad) and imaged on a ChemiDoc MP Imaging System (Bio-Rad).

***Preparation of M13 phage particles displaying the anti-GFP intrabody***

5-alpha F' *E. coli* cells (New England Biolabs, C2992H) harboring the Sec-anti-GFP-ib-p3c phagemid (**table S1**) were grown in 2YT medium supplemented with tetracycline (15 µg/mL) and carbenicillin (50 µg/mL) containing 3% glucose at 37 °C until reaching an optical density at 600 nm of 0.5. M13KO7 helper phage (New England Biolabs, N0315S) was then added at a multiplicity of infection of 10. Following infection, cultures were incubated at 37 °C for 30 min. Cells were collected by centrifugation at 5,000 × g for 5 min at room temperature and resuspended in fresh 2YT medium supplemented with carbenicillin (50 µg/mL) and kanamycin (50 µg/mL), but lacking glucose, to induce expression of the anti-GFP-ib-p3c fusion protein. Phage production was allowed to proceed overnight at 30 °C. Phage-containing supernatants were collected by centrifugation at 17,000 × g to pellet bacterial cells. Phage particles were precipitated by adding polyethylene glycol (PEG 8000)/2.5 M NaCl solution to the clarified supernatant at a 1:5 volume ratio, followed by thorough mixing and incubation on ice for 1 h. Precipitated phage particles were collected by centrifugation at 17,000 × g at 4 °C and resuspended in PBS at 1% of the original culture volume. Phage titers were determined spectrophotometrically using the following equation: Phage titer (cfu/mL) = $[(A_{269} - A_{320}) \times 6 \times 10^{16}]$ / phagemid size (bp) (*48*). For long-term storage, phage preparations were supplemented with glycerol to a final concentration of 14% and stored at -20 °C.

***Basic image processing, particle detection, and tracking***

Raw fluorescence microscopy movies were imported into MicroLive (*47*) together with acquisition metadata, including pixel size and frame interval. Image data were processed as multidimensional time-lapse stacks comprising three spatial dimensions (x, y, and z), two fluorescence channels corresponding to translation and folding, and time. Movies were included for analysis only if cells remained morphologically healthy, stayed within the focal range throughout acquisition, and contained clearly detectable translation spots.

Cell regions were segmented either automatically using a watershed-based algorithm or manually by tracing cell boundaries. Segmentation masks were visually inspected prior to downstream analysis, and regions containing large intracellular aggregates or segmentation artifacts were excluded. Photobleaching correction was performed by calculating the median fluorescence intensity of each frame across the entire field of view and fitting the resulting decay profile to an exponential function.

For particle detection and tracking, z-stacks were first converted to maximum-intensity projections at each time point. Translation spots were then detected and linked over time in MicroLive (*47*) using an expected spot diameter of 5 pixels, a linking radius of 10 pixels, and a one-frame gap allowance to accommodate transient missed detections. Spot intensities were quantified following local background correction by subtracting the fluorescence intensity of the surrounding region from that measured within the particle region (*49*).

***Harringtonine run-off assay analysis***

For harringtonine run-off assays, stable cells expressing anti-HA-ib-Halo, anti-GFP-ib-mSG, and tdMCP-CAAX, or parental cells, were seeded in MatTek chambers. Stable cells were transiently transfected with 2.5 μg of the indicated coTFT reporter plasmid, whereas parental cells were co-transfected with CMV-anti-HA-ib-Halo (table S1), CMV-tdMCP-CAAX (table S1), and the indicated coTFT reporter plasmid. Live-cell imaging was performed 6-10 h post-transfection.

Image processing and quantification were carried out in MicroLive (*47*). Cells were segmented prior to analysis, and no photobleaching correction was applied because of the low temporal sampling frequency. Translation spots were detected in each frame using a user-defined detection threshold, and spot intensities were quantified over time. To avoid artificially inflating the average spot intensity as the number of detectable spots declined during run-off, post-treatment intensities were normalized to the average pre-treatment spot count rather than the instantaneous spot count. The resulting intensity vector $I(t)$, representing the average signal per pre-treatment translation spots, was defined as:

$$I(t) = \frac{1}{N(t)} \sum_{i=1}^{n(t)} I_i(t),$$

where

$$N(t) = \begin{cases} n(t), & t < t_H \\ \langle n \rangle_{t \leq t_H}, & t \geq t_H \end{cases},$$

and $\langle n \rangle$ denotes the average number of translation spots, $n(t)$, across all frames preceding harringtonine addition ($t_H$).

The reported population-averaged normalized intensity was background-corrected by subtracting a baseline intensity defined as the mean of the last 10 frames in the analysis window and then rescaled to its pre-treatment mean. This normalization scales the intensity trajectory from approximately 1 before harringtonine treatment to ~0 at the post-run-off baseline.

To estimate translation elongation rates, a linear function was fitted to the population-averaged normalized intensity decay over its linear phase after harringtonine addition. The fitting window excluded the late, slowly decaying tail of the curve, which was attributed to elongation heterogeneity and ribosome stalling. Population means and standard errors of the mean (SEMs) were calculated across all responding cells at each time point, and the resulting mean trajectory was fitted by linear regression. The run-off time was defined as the time at which the extrapolated linear fit intersected the baseline, corresponding to complete clearance of translating ribosomes from the reporter mRNA. Translation elongation rates were then estimated from the run-off time and the effective fluorescent length of the reporter:

$$k_e = \frac{L_{eff}}{T_{runoff}},$$

where $L_{eff} = L_1 + L_2/2$ is the effective fluorescent run-off length. Here $L_1$ is the length of the coding sequence downstream of the smHA tag, and $L_2$ is the length of the smHA tag. For the 4×dGFP$_{Fast}$ and 4×dGFP$_{Slow}$ reporters ($L_1$ = 1,491 aa; $L_2$ = 335 aa), $L_{eff}$ = 1,659 aa. For the XBP1u containing reporters, $L_{eff}$ was 1,685 aa for 4×dGFP$_{Fast}$-XBP1u and 1,688 aa for XBP1u-4×dGFP$_{Fast}$. To account for the delay associated with harringtonine diffusion and inhibition of translation initiation, 1 min was subtracted from the fitted run-off time ($T_{runoff}$) before calculating the elongation rate.

***Puromycin assay analysis***

For puromycin assays, stable cells expressing anti-HA-ib-Halo and anti-GFP-ib-mSG were seeded in MatTek chambers and transiently transfected with 2.5 µg of the indicated coTFT reporter plasmid. Live-cell imaging was performed 6 h after transfection. Image processing was implemented using MicroLive (*47*). First, cells were segmented, regions containing visible protein aggregates were manually excluded from analysis. Translation spots were then detected in each frame using a particle size of 5 pixels and a user-defined intensity threshold. Unlike the harringtonine run-off assay, which quantified the gradual decay of spot fluorescence intensity, puromycin assays used translation spots count as the primary quantitative readout. Spot counts were quantified at each frame and normalized to the average pre-treatment spot count for each cell to account for cell-to-cell variability. Normalized spot counts were then averaged across all analyzed cells, and results are reported as means and SEMs.

***Quantification of co-translational folding efficiency***

For co-translational folding efficiency quantification, stable cells expressing anti-HA-ib-Halo and anti-GFP-ib-mSG were seeded in MatTek chambers and transiently transfected with 2.5 µg of the indicated coTFT reporter plasmid. The cells were imaged 6-10 h post-transfection. Quantification was carried out in MicroLive (*47*). First, cells were segmented, regions containing visible protein aggregates were manually excluded from analysis. Then, translation spots were detected in the nascent-chain channel using a particle size of 5 pixels and tracked using a linking radius of 7 pixels. To exclude spurious detections, only spots persisting for at least three consecutive frames were retained for analysis. For each detected translation spot, the corresponding coordinates were used to extract an averaged image crop for the folding channel. These image crops were subsequently classified by the convolutional neural network (CNN) in MicroLive (*47*) for the presence (colocalized translation and folding spots) or absence of a folding spot. Automated classifications were then reviewed manually, and each crop was visually inspected to validate the assigned classification. Co-translational folding efficiency, $c_{eff}$, was calculated for each cell as:

$$c_{eff} = \left(\frac{n_{cs}}{n_{ts}}\right) \times 100,$$

where $n_{cs}$ is the number of colocalized translation and folding spots in a cell, and $n_{ts}$ is the total number of detected translation spots in the cell. Population means and SEMs of co-translation folding efficiency were calculated across all analyzed cells.

#### *Modeling approach for folding time estimation*

To estimate the folding times of $\text{dGFP}_{\text{Fast}}$ and $\text{dGFP}_{\text{Slow}}$ on translating ribosomes, we developed a mathematical model linking translation elongation to protein folding. The model is based on a spatial-constraint framework in which protein domains must first be translated and emerge from the ribosome exit tunnel before folding can occur. Co-translational folding is therefore limited by both the time required for folding and the remaining translation time before termination.

The reporter transcript encodes an open reading frame (ORF) of length $L_{total}$ = 1,826 aa, consisting of an N-terminal smHA tag (345 aa, including an 11-aa linker), up to six tandem dGFP domains (245 aa each), and an 11-aa C-terminal region. A dGFP domain is fully synthesized when the ribosome reaches its terminal residue at position $x_{end}$. An additional 50 aa ($\ell_{tunnel}$) elongation is required before the entire dGFP domain emerges from the ribosome exit tunnel and becomes competent to fold (*50*). The emergence position of each dGFP domain was therefore defined as:

$$x_{exit} = min\ (x_{end} + \ell_{tunnel}, L_{total}).$$

The model incorporates four assumptions. First, ribosomes elongate at a constant rate $k_e$ (aa/s), determined experimentally from harringtonine run-off assays of the 4×$\text{dGFP}_{\text{Fast}}$ and 4×$\text{dGFP}_{\text{Slow}}$ reporters. Second, each dGFP domain is characterized by a folding time $\tau_{fold}$. At a constant elongation rate, this folding time corresponds to an effective translation distance $L_{fold}$:

$$L_{fold} = \tau_{fold} \times k_e.$$

Third, a dGFP domain becomes folded and is recognized by the fluorescent anti-GFP intrabody only after the ribosome has translated an additional $L_{fold}$ amino acids beyond its emergence position. The fluorescence onset position for domain $(i)$ is given by

$$x_{fluorescent}^{(i)} = x_{exit}^{(i)} + L_{fold}.$$

Fourth, assuming a uniform ribosome density under steady-state translation, the probability that dGFP domain $(i)$ contributes to co-translational folding efficiency measurement is proportional to the fraction of the encoding sequence remaining after its fluorescence onset position:

$$P_i = f(x) = \begin{cases} \frac{L_{total} - x_{fluorescent}^{(i)}}{L_{total}}, & x_{fluorescent}^{(i)} < L_{total} \\ 0, & otherwise \end{cases}.$$

The predicted co-translational folding efficiency for a reporter containing $n$ dGFP domains is:

$$E(n) = A \sum_{i=1}^{n} P_i,$$

where $A$ is a scaling factor that accounts for dGFP copy-number-independent effects (e.g., intrabody binding efficiency, productive folding yield, and detection sensitivity) on co-translational folding efficiency.

The model contains two fitted parameters: the folding time $\tau_{fold}$ and the scaling factor $A$. These parameters were estimated by fitting the model to the experimentally measured co-translational folding efficiencies of the 1× - 6×dGFP reporters (**Figs. 4C-D**) through minimization of $\chi^2$ using SEM-weighted residuals. For $dGFP_{Fast}$ ($k_e$ = 3.88 aa/s), the best-fit parameters were $\tau_{fold}$ = 88.8 s [95% CI: 51.0 - 114.3 s] and $A$ = 66.5% [95% CI: 51.3 - 82.9%], corresponding to $L_{fold}$ = 345 aa. For $dGFP_{Slow}$ ($k_e$ = 1.63 aa/s), the best-fit parameters were $\tau_{fold}$ = 332.3 s [95% CI: 238.4 - 408.0 s] and $A$ = 38.9% [95% CI: 28.1 - 54.8%], corresponding to $L_{fold}$ = 542 aa. Parameter identifiability was assessed by evaluating $\chi^2$ values across a two-dimensional grid of $\tau_{fold}$ and $A$ (**fig. S6**). Spatial cutoffs for individual dGFP domains are summarized in **table S2**, and the corresponding model fits are shown in **Figs. 4E** and **4F**.

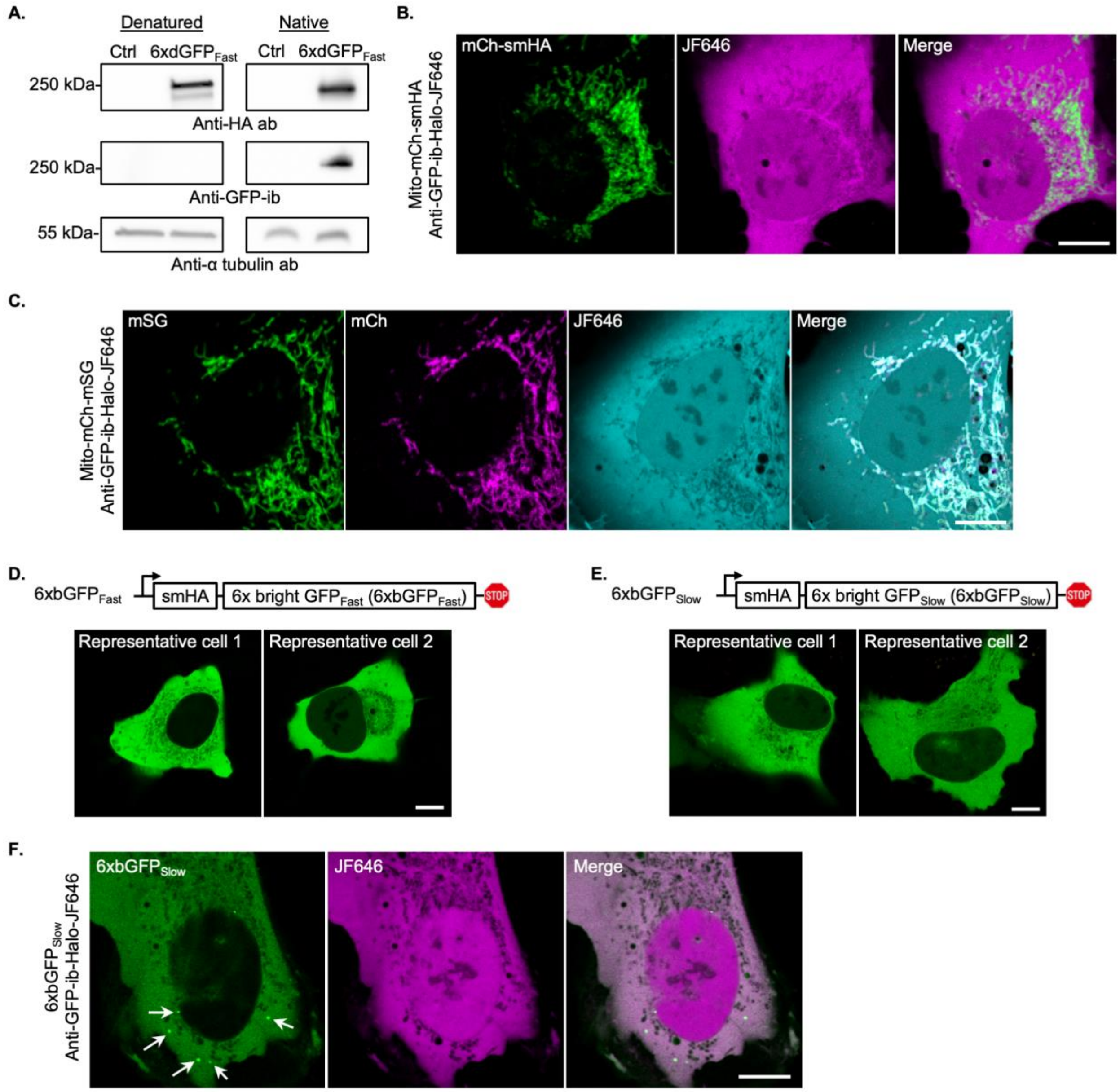


**Fig. S1. Validation of the co-translational folding tracking technology. (A)** Representative western blot showing that anti-GFP-ib binds the native folded 6×dGFP$_{\text{Fast}}$ reporter protein (right) but not the denatured protein (Left), confirming conformation-dependent recognition. The anti-HA antibody blot verifies full-length reporter expression and α-tubulin serves as a loading control. **(B)** Representative cell co-expressing Mito-mCh-smHA and HaloTag fused anti-GFP-ib stained with JF646 (anti-GFP-ib-Halo-JF646), demonstrating that anti-GFP-ib does not bind smHA or mCh. **(C)** Representative cell co-expressing Mito-mCh-mSG and anti-GFP-ib-Halo-JF646, demonstrating that anti-GFP-ib does not bind mCh or mSG. **(D-E)** Representative images of cells expressing the bright reporters $6\times\text{bGFP}_{\text{Fast}}$ (D) and $6\times\text{bGFP}_{\text{Slow}}$ (E) 24 h after transfection. Robust green fluorescence confirms that the GFP repeats in both reporters fold properly and remain fluorescent without substantial aggregation. **(F)** Representative cell co-expressing $6\times\text{bGFP}_{\text{Slow}}$ and anti-GFP-ib-Halo-JF646, demonstrating that anti-GFP-ib does not recognize aggregated reporter protein (white arrows). Scale bars, 10 µm.

```
A.
dGFP_Fast  1    SKGEELFTGVVPILVELDGDVNGHKFSVRGEGEGDATNGKLTLKFICTTGKLPVPWPTLV  60
dGFP_Slow  1    ............................S........Y......................  60

dGFP_Fast  61   TTLTYAVQCFSRYPDHMKRHDFFKSAMPEGYVQERTISFKDDGTYKTRAEVKFEGDTLVN  120
dGFP_Slow  61   ..FS.......................................N................  120

dGFP_Fast  121  RIELKGIDFKEDGNILGHKLEYNFNSHNVYITADKQKNGIKANFKIRHNVEDGSVQLADH  180
dGFP_Slow  121  .......................Y.........................I..........  180

dGFP_Fast  181  YQQNTPIGDGPVLLPDNHYLSTQSVLSKDPNEKRDHMVLLEFVTAAGITHGMDELYK     237
dGFP_Slow  181  ........................K................................     237

B.
dGFP_Fast  1    AGCAAGGGCGAGGAGCTGTTCACCGGGGTGGTGCCCATCCTGGTCGAGCTGGACGGCGAC  60
dGFP_Slow  1    ............................................................  60

dGFP_Fast  61   GTAAACGGCCACAAGTTCAGCGTGCGCGGCGAGGGCGAGGGCGATGCCACCAACGGCAAG  120
dGFP_Slow  61   ........................A..........................T........  120

dGFP_Fast  121  CTGACCCTGAAGTTCATCTGCACCACCGGCAAGCTGCCCGTGCCCTGGCCCACCCTCGTG  180
dGFP_Slow  121  ............................................................  180

dGFP_Fast  181  ACCACCCTGACCTACGCTGTGCAGTGCTTCAGCCGCTACCCCGACCACATGAAGCGCCAC  240
dGFP_Slow  181  ......T.T.G.................................................  240

dGFP_Fast  241  GACTTCTTCAAGTCCGCCATGCCCGAAGGCTACGTCCAGGAGCGCACCATCAGCTTCAAG  300
dGFP_Slow  241  ............................................................  300

dGFP_Fast  301  GACGACGGCACCTACAAGACCCGCGCCGAGGTGAAGTTCGAGGGCGACACCCTGGTGAAC  360
dGFP_Slow  301  ..........A.................................................  360

dGFP_Fast  361  CGCATCGAGCTGAAGGGCATCGACTTCAAGGAGGACGGCAACATCCTGGGGCACAAGCTG  420
dGFP_Slow  361  ............................................................  420

dGFP_Fast  421  GAGTACAACTTCAACAGCCACAACGTCTATATCACCGCCGACAAGCAGAAGAACGGCATC  480
dGFP_Slow  421  ..........A.................................................  480

dGFP_Fast  481  AAGGCCAACTTCAAGATCCGCCACAACGTGGAGGACGGCAGCGTGCAGCTCGCCGACCAC  540
dGFP_Slow  481  ...........................A.T..............................  540

dGFP_Fast  541  TACCAGCAGAACACCCCCATCGGCGACGGCCCCGTGCTGCTGCCCGACAACCACTACCTG  600
dGFP_Slow  541  ............................................................  600

dGFP_Fast  601  AGCACCCAGTCCGTGCTGAGCAAAGACCCCAACGAGAAGCGCGATCACATGGTCCTGCTG  660
dGFP_Slow  601  ............AA..............................................  660

dGFP_Fast  661  GAGTTCGTGACCGCCGCCGGGATCACTCACGGCATGGACGAGCTGTACAAG  711
dGFP_Slow  661  ...................................................  711
```

**Fig. S2. Comparison of protein and nucleotide sequences between dGFP$_{Fast}$ and dGFP$_{Slow}$.** **(A)** Protein sequence alignment. **(B)** Nucleotide sequence alignment. Differences between the two sequences are highlighted in bold, whereas dots denote identical amino acids or nucleotides.

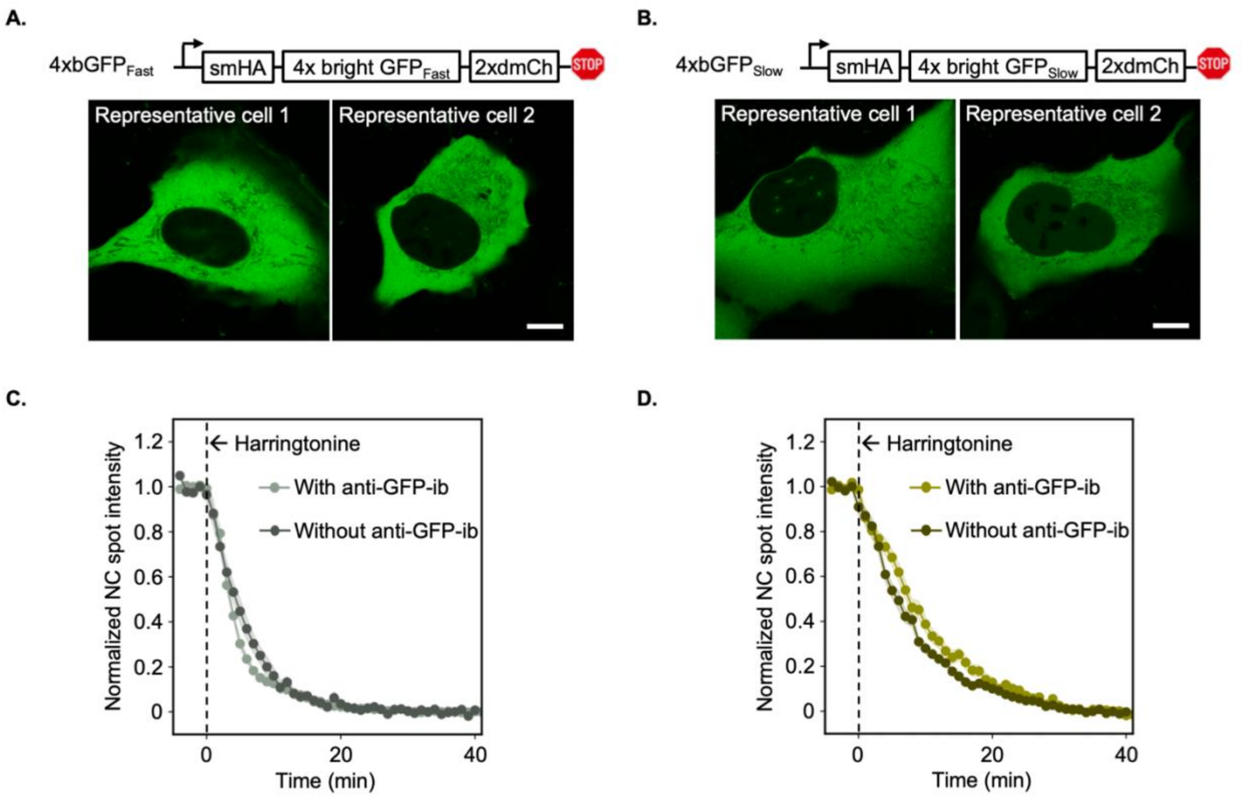


**Fig. S3. Fast- and slow-folding reporters exhibit corresponding differences in translation elongation rates. (A-B)** Representative images of cells expressing $4\times bGFP_{Fast}$ (A) or $4\times bGFP_{Slow}$ (B) 24 h after transfection. **(C-D)** Harringtonine run-off assay performed in the presence or absence of the anti-GFP intrabody for $4\times dGFP_{Fast}$ (C; n = 12 cells) and $4\times dGFP_{Slow}$ (D; n = 15 cells). Curves obtained in the presence of the anti-GFP intrabody were replotted from Figs. 2H-J for comparison. Scale bars, 10 µm.

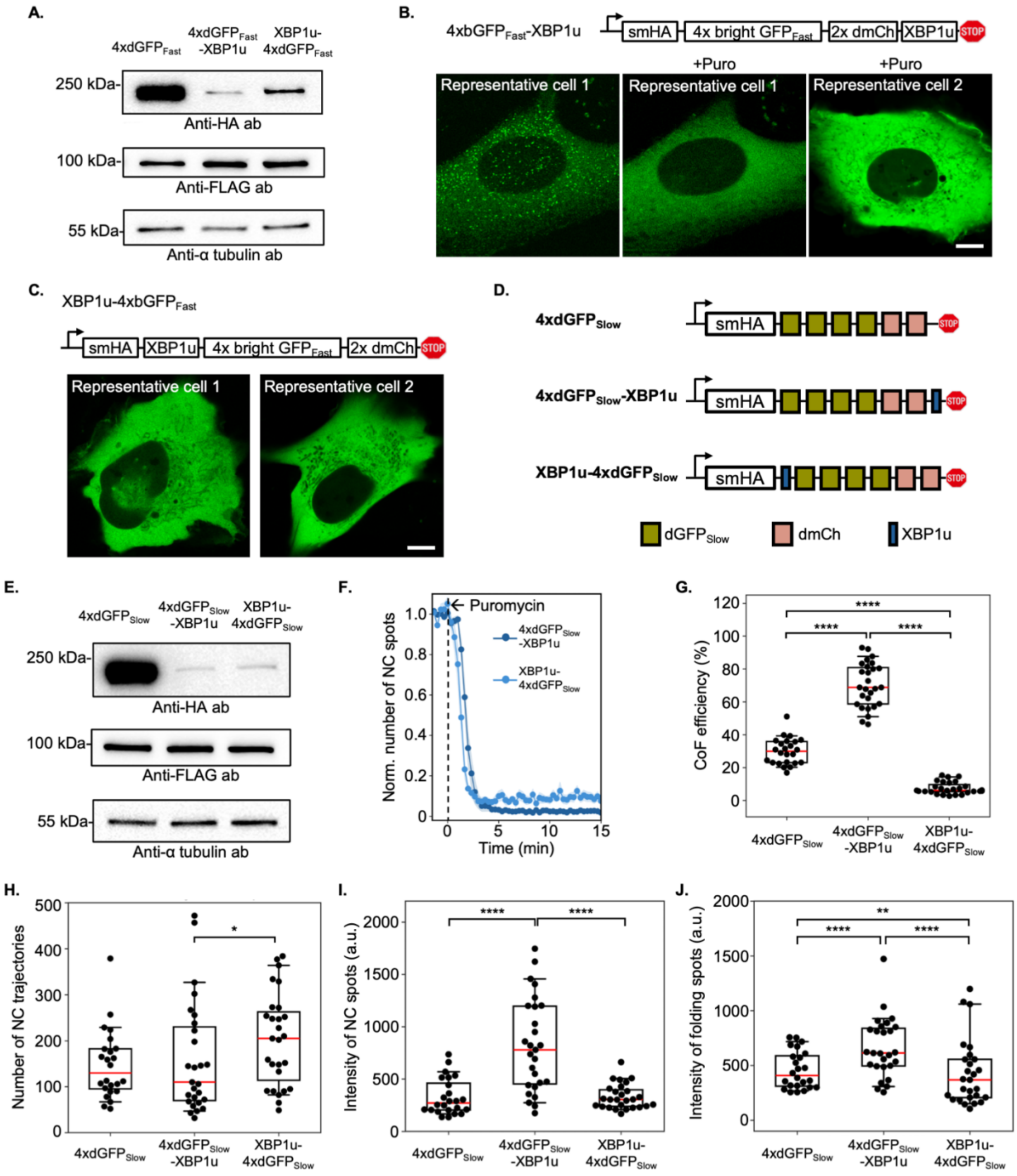
A.
4xdGFPFast
4xdGFPFast -XBP1u
XBP1u-4xdGFPFast
250 kDa-
Anti-HA ab
100 kDa-
Anti-FLAG ab
55 kDa-
Anti-α tubulin ab
B.
4xbGFPFast-XBP1u
smHA
4x bright GFPFast
2x dmCh
XBP1u
STOP
+Puro
+Puro
Representative cell 1
Representative cell 1
Representative cell 2
C.
XBP1u-4xbGFPFast
smHA
XBP1u
4x bright GFPFast
2x dmCh
STOP
Representative cell 1
Representative cell 2
D.
4xdGFPSlow
4xdGFPSlow-XBP1u
XBP1u-4xdGFPSlow
smHA
STOP
dGFPSlow
dmCh
XBP1u
E.
4xdGFPSlow
4xdGFPSlow -XBP1u
XBP1u-4xdGFPSlow
250 kDa-
Anti-HA ab
100 kDa-
Anti-FLAG ab
55 kDa-
Anti-α tubulin ab
F.
Puromycin
4xdGFPSlow -XBP1u
XBP1u-4xdGFPSlow
Norm. number of NC spots
Time (min)
G.
CoF efficiency (%)
4xdGFPSlow
4xdGFPSlow -XBP1u
XBP1u-4xdGFPSlow
H.
Number of NC trajectories
I.
Intensity of NC spots (a.u.)
J.
Intensity of folding spots (a.u.)

349

**Fig. S4. Perturbing translation elongation alters co-translation folding outcomes. (A)** Representative western blot of the 4×dGFP$_{Fast}$ reporters with or without XBP1u expressed for 24 h, confirming full-length protein expression. 4×FLAG-mCh-β-actin was used as a transfection control and α-tubulin as a loading control. **(B)** Representative images of cells expressing the 4×bGFP$_{Fast}$-XBP1u reporter 24 h after transfection. Robust green fluorescence confirms proper folding of the GFP repeats in the reporter. Puromycin-responsive puncta (representative cell 1) are bona fide nascent chain spots illuminated by folded and fluorescent GFP repeats in the bright reporter, consistent with prolonged translation pause induced by XBP1u. **(C)** Representative images of cells expressing the XBP1u-4×bGFP$_{Fast}$ reporter 24 h after transfection, confirming proper folding of the GFP repeats. **(D)** Schematic of the 4×dGFP$_{Slow}$ reporters with or without XBP1u. **(E)** Representative western blot of the 4×GFP$_{Slow}$ reporters with or without XBP1u expressed for 24 h, confirming full-length protein expression. 4×FLAG-mCh-β-actin was used as a transfection control and α-tubulin as a loading control. **(F)** Puromycin assay results for 4×dGFP$_{Slow}$-XBP1u ($n = 8$ cells) and XBP1u-4×dGFP$_{Slow}$ ($n = 8$ cells). **(G)** Box-and-whisker plots of co-translational folding (coF) efficiencies for 4×dGFP$_{Slow}$ (29.67 ± 1.61%, $n = 24$ cells, 3,538 trajectories), 4×dGFP$_{Slow}$-XBP1u (70.04 ± 2.61%, $n = 27$ cells, 4,315 trajectories), and XBP1u-4×dGFP$_{Slow}$ (7.47 ± 0.71%, $n = 27$ cells, 5,482 trajectories). **(H)** Box-and-whisker plots of the number of nascent chain (NC) trajectories per cell corresponding to the data in (G). **(I)** Box-and-whisker plots of the median NC intensity per cell corresponding to the data in (G). **(J)** Box-and-whisker plots of the median folding-spot intensity per cell for 4×dGFP$_{Slow}$ (1,096 trajectories), 4×dGFP$_{Slow}$-XBP1u (3,257 trajectories), and XBP1u-4×dGFP$_{Slow}$ (383 trajectories) corresponding to the data in (G). In box-and-whisker plots, the red line indicates the median. *$p < 0.05$, **$p < 0.01$, ****$p < 0.0001$ (Mann-Whitney U test). Scale bars, 10 μm.

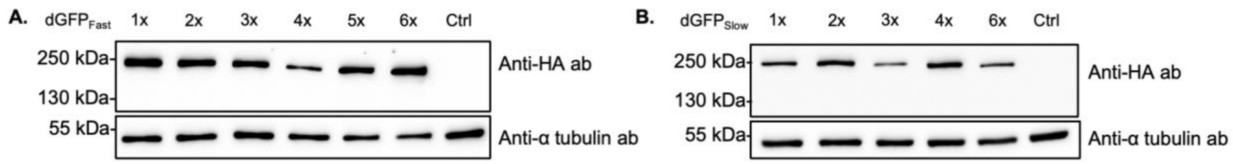


**Fig. S5. Western blot analysis of the dGFP reporter series used for folding time prediction.** Representative western blots of the 1× - 6×dGFP$_{Fast}$ **(A)** and 1× - 6×dGFP$_{Slow}$ **(B)** reporter series used for folding time prediction, confirming full-length protein expression of all reporters. α-Tubulin served as a loading control.

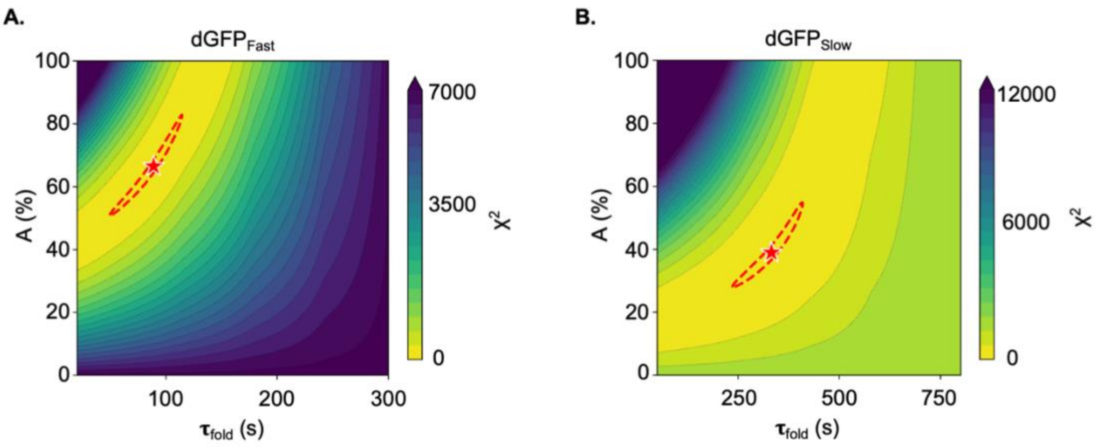


**Fig. S6. Parameter sensitivity analysis.** Two-dimensional $\chi^2$ landscape over folding time ($\tau_{fold}$) and scaling factor ($A$) for dGFP$_{Fast}$ **(A)** and dGFP$_{Slow}$ **(B)**. Color scale indicates $\chi^2$ value, clipped at the 95th percentile. Red stars denote the best-fit parameter values, and red dashed contours indicate the 95% confidence regions.

**Table S2. Spatial cutoff analysis for individual dGFP domains in the coTFT reporters**

| | | | $6\times dGFP_{Fast}$ | | $6\times dGFP_{Slow}$ | |
|---|---|---|---|---|---|---|
| Domain | $x_{end}$ (aa) | $x_{exit}$ (aa) | $x_{fluorescent}$ (aa) | Status | $x_{fluorescent}$ (aa) | Status |
| dGFP 1 | 590 | 640 | 985 | Folds | 1182 | Folds |
| dGFP 2 | 835 | 885 | 1230 | Folds | 1427 | Folds |
| dGFP 3 | 1080 | 1130 | 1475 | Folds | 1672 | Folds |
| dGFP 4 | 1325 | 1375 | 1720 | Folds | 1917 | Fails |
| dGFP 5 | 1570 | 1620 | 1965 | Fails | 2162 | Fails |
| dGFP 6 | 1815 | 1826 | 2171 | Fails | 2368 | Fails |
